# MuCST: restoring and integrating heterogeneous morphology images and spatial transcriptomics data with contrastive learning

**DOI:** 10.1101/2024.06.26.600708

**Authors:** Yu Wang, Xiaoke Ma

**Affiliations:** School of Computer Science and Technology, Xidian University, No.2 South Taibai Road, Xi’an, 710071, Shaanxi, China; Key Laboratory of Smart Human-Computer Interaction and Wearable Technology of Shaanxi Province, Xidian University, No.2 South Taibai Road, Xi’an, 710071, Shaanxi, China

## Abstract

Spatially resolved transcriptomics simultaneously measure the spatial location, histology images, and transcriptional profiles of the same cells or regions in undissociated tissues. Integrative analysis of multi-modal spatially resolved data holds immense potential for understanding the mechanisms of biology. Here we present a flexible multi-modal contrastive learning for the integration of spatially resolved transcriptomics (MuCST), which jointly perform denoising, elimination of heterogeneity, and compatible feature learning. We demonstrate that MuCST robustly and accurately identifies tissue subpopulations from simulated data with various types of perturbations. In cancer-related tissues, MuCST precisely identifies tumor-associated domains, reveals gene biomarkers for tumor regions, and exposes intra-tumoral heterogeneity. We also validate that MuCST is applicable to diverse datasets generated from various platforms, such as STARmap, Visium, and omsFISH for spatial transcriptomics, and hematoxylin and eosin or fluorescence microscopy for images. Overall, MuCST not only facilitates the integration of multi-modal spatially resolved data, but also serves as pre-processing for data restoration (Python software is available at https://github.com/xkmaxidian/MuCST).

## Introduction

Cells are the fundamental units of tissues in multicellular organisms, where cells are organized into groups of similar cells physically clustered together with various states. Recognizing the structure and spatial location of cells is vital for understanding the emergent properties and pathology of tissues (1). The traditional microscopy technology identifies and characterizes cell groups (also called cell types or cell sub-populations) through similarities of morphology, including shapes, sizes and physical appearance of cells (2). However, morphology alone is insufficient to fully characterize structure of cells because of the unstable states of cells (3). Fortunately, the single-cell RNA sequencing (scRNA-seq) technology enables generation of whole genome-wide expression at cell level, providing complementary information to characterize structure of cells at molecular level (4,5).

However, the dissociation step of scRNA-seq erases spatial context of cells from their original tissues that is crucial for understanding cellular functions and organizations (6). Spatial Transcriptomics (ST) (7) simultaneously allows morphological and transcriptional profiling of cells in the same tissue regions, which also retains spatial context of cells (8). Typically, current ST technologies can be broadly divided into two categories, i.e., imaging- and next-generation sequencing (NGS)-based methods, where the former one uses probes to localize mRNA transcripts, including FISH and MERFISH (9), seqFISH (10), and STARmap (11), which are criticized for their limited capacity to detect RNA transcripts. To overcome this limitation, The latter one utilizes spatial barcode and next-generation sequencing to retain transcription and spatial information, including Legacy Spatial Transcriptomics (12), 10x Visium (13), Slide-seq (14) and Stereo-seq (15). The accumulated spatially resolved data provides a great opportunity to investigate functions and cellular structure of tissues by exploiting interesting patterns and features that cannot be discerned from scRNA-seq data (16).

Therefore, integrative analysis of spatially resolved data is a prominent task since it sheds light on revealing mechanisms of tissues. On the basis of principles of algorithms, current approaches are roughly divided into two categories, i.e., transcript- and image-based methods, where the former ones are devoted to integrate transcriptomics and spatial information, and the latter ones fuse morphology, transcript, and spatial information. Specifically, transcript-based approaches concentrate on learning cell features by balancing transcriptomics and spatial coordinates of cells with various strategies. For example, algorithms for scRNA-seq data, such as SCANPY (17), DRjCC (18) and jSRC (19) are directly applied to ST data by ignoring spatial information of cells, resulting in the undesirable performance. To overcome this limitation, many algorithms are developed by incorporating spatial coordinates into feature learning with various manners. For instance, Gitto (20) employs the hidden Markov random field model, whereas BayesSpace (21) adopts the Bayesian statistical method. STAGATE (22) and GraphST (23) utilize graph neural networks, while SEDR (24), DRSC (25), and SpatialPCA (26) learns the low-dimensional features of cells in subspaces with dimension reduction. Spatial-MGCN (27) constructs neighbor graphs with transcriptional feature and spatial information, and employs graph neural network to learn features of cells.

Nevertheless, these algorithms ignore morphological information in histology images that usually provide vital supplementary information for transcriptomics. For example, transcriptional variations within distinct spatial domains are often mirrored in morphology (29). However, integrating morphology and spatial transcriptomics is highly non-trivial because of the extra-ordinary heterogeneity of data. Current algorithms leverage morphological information in histology images to complement spatial transcriptomics with different strategies. For example, stLearn (28) calculates morphological distance between cells to smooth and augment expression of cells. SpaGCN (29) and DeepST (30) integrate spatial and morphological information into cell networks and learns features with graph convolution network (GCN). stMVC (31) and stMGATF (32) transform spatially resolved transcriptomics into multi-view clustering and then adopt semi-supervised strategy for down-stream analysis. MUSE (33) integrates information of morphology and transcription to learn joint representation with deep learning, while ConGI (34) executes integrative analysis with multi-modality contrastive learning, and conST (35) constructs attributed cell networks to obtain features with contrastive learning by exploiting topological structure of networks.

Even though few attempt have been devoted to the integration of histology images and spatial transcriptomics, there are still many unsolved and critical problems. First, the extra experimental steps required to preserve the locations of sequencing results in noise in spatially resolved data (36), which poses a great challenge in designing effective integrative algorithms. Current algorithms remove noise in the pre-processing procedure, which separates noise and feature learning, failing to fully characterize and model noise. Second, spatially resolved data are highly heterogeneous, and current algorithms directly learn the low-dimensional features of cells, which neglects the heterogeneity of spatial omic data. How to avoid heterogeneity of spatial data is still a great challenge for integrative analysis. Third, spatially resolved data consist of multiple modalities that are complementary to each other, and current algorithms fail to deeply fuse all modalities, thereby reducing the quality of cell features. How to learn compatible cell features is also vital for integrating spatially resolved data.

To address the aforementioned issues, we present a novel and flexible <u>Mu</u>ltimodal <u>C</u>ontrastive learning for the integration of <u>S</u>patially resolved <u>T</u>ranscriptomics (MuCST), including morphology, spatial coordinates and transcription profiles of cells. As shown in Fig. 1, MuCST consists of two major components, i.e., cell feature learning and data restoration, where the former procedure obtains heterogeneous morphological and transcriptional features of cells, and data restoration removes heterogeneity and noise of features with multi-modal contrastive learning. Therefore, MuCST simultaneously joins feature learning, denoising, and elimination of heterogeneity of spatially resolved transcriptomics into an overall framework, which significantly enhances discriminative and compatibility of cell features. The experimental results on the simulated and spatially resolved transcriptomics data demonstrate that MuCST not only precisely removes noise, but also improves the dissection and interpretation of spatial heterogeneity of tissues.

**Fig. 1.**
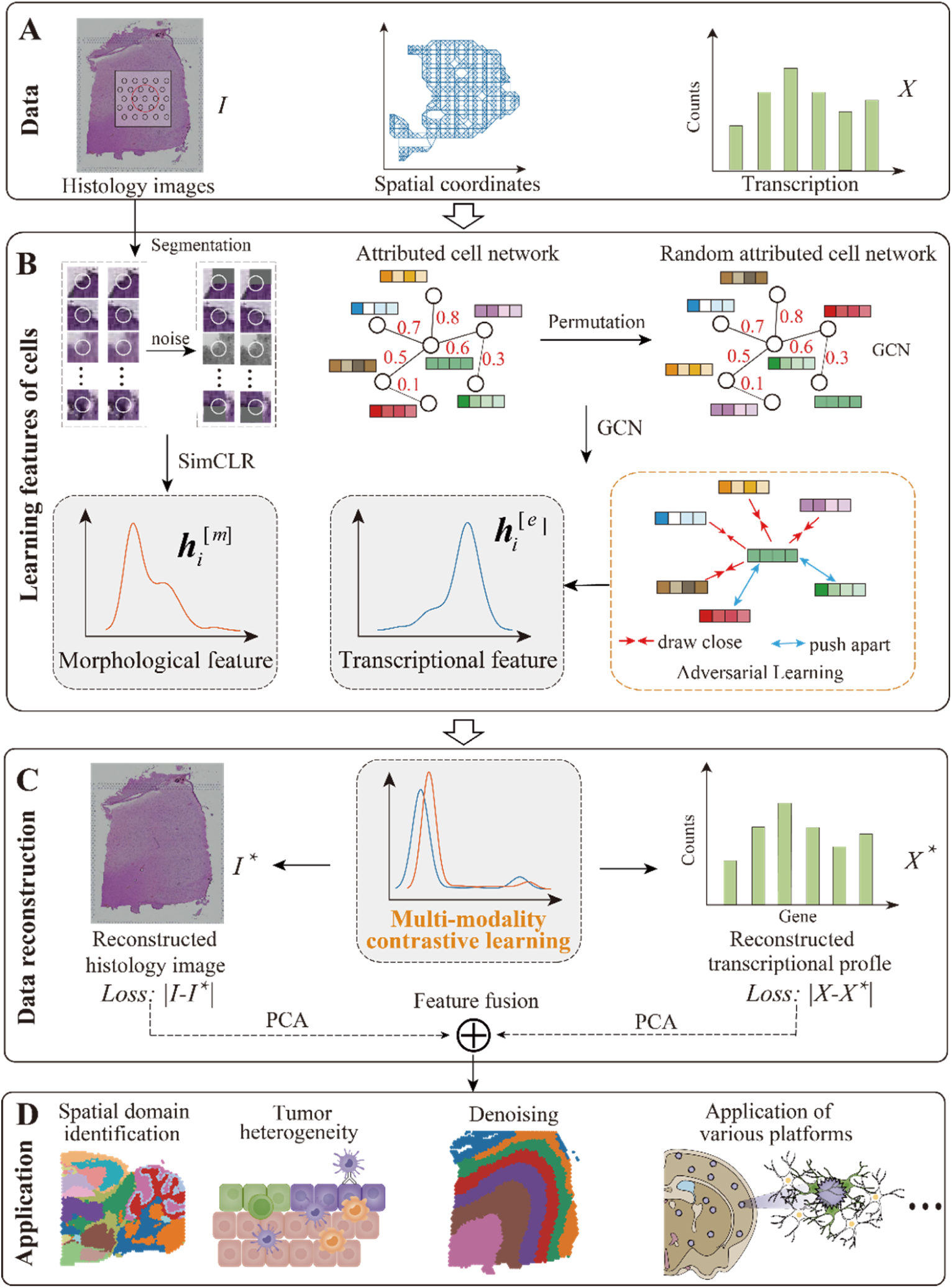
Overview of MuCST for integrating spatially resolved data with histology images, spatial coordinates and transcriptional information. **(A)** Spatially resolved multi-modal data include histology images, spatial coordinates of cells, and transcriptional profiles of cells. **(B)** MuCST learns the morphological feature of cells from histology images with available SimCLR, and learns the transcriptional features of cells by integrating spatial and transcriptional information, where an attributed cell network model is proposed. Graph convolution network (GCN) is employed to learn compatible transcriptional features with adversarial learning. **(C)** MuCST reconstructs histology images and expression profiles of cells with consistent cell features that are obtained with multi-modal contrast learning by fusing the morphological and transcriptional features of cells. **(D)** Down-stream analysis of fused cell features from the reconstructed image and expression profiles with principle component analysis (PCA) include spatial domain identification, tumor micro-environment, spatial transcriptomics data denoising, and so on.

## Results

### Overview of MuCST

To facilitate the understanding of this study, the rationale and procedures of MuCST are first presented in this section (Technical details can be referred to section of methods). For clarity, we utilize “cell” to denote the basic measurement units for imaging-based ST technologies, and “spot” to denote the basic measurement units for barcode-based ST technologies. We explain the workflow of MuCST using barcode-based spatially resolved transcriptomics data as an example for convenience.

Spatially resolved transcriptomics data comprehensively cover histology images, spatial coordinates and transcription of spots (Fig. 1**A**), posing a great challenge on integrative analysis of them because of extra-ordinary heterogeneity of multi-modal data. Available algorithms directly fuse various types of spots’ features, failing to appropriately address heterogeneity and intrinsic structure of data, resulting in undesirable performance. To address these issues, we propose a novel and flexible MuCST algorithm for the integration of histological images and spatial transcriptomics with a hierarchical structure, which consists of three major components, i.e., learning the morphological and transcriptional features of spots, data reconstruction with multimodal contrast learning, and downstream analysis (Fig. 1). The underlying assumption is that spatial resolved data characterize tissues from different perspectives and levels, and integrative analysis of these data with a refined ordering according to their roles is promising for heterogeneous data. Specifically, spatial and expression profiles of spots characterize tissues from micro-level, whereas histological images depict tissues from macro-level. Thus, MuCST first integrates spatial coordinates and transcriptomics to obtain transcriptional features of spots, and then fuses morphological and transcriptional features of spots with multi-modal contrast learning.

On the feature learning issue, MuCST independently learns the morphological and transcriptional features of spots to avoid heterogeneity of data learning issue (Fig. 1**B**). Specifically, MuCST splits the morphological image I into patches for each spot, and these patches are randomly noised. And, the pre-trained DNN model (38), followed by multi-layer perception (MLP) (39) are adopted to learn the latent morphological features of spots. To enhance quality of features, SimCLR (37) is employed to discriminate the original and noised patches (Fig. 1**B**). And, MuCST learns the transcriptional features of spots by proposing a network-based model with graph convolution network (GCN), where the indirect topological structure is exploited to fully characterize spatial and transcriptional information of spatially resolved data. Specifically, MuCST first constructs an attributed network by integrating histological images, spatial coordinates and expression profiles *X* of spots (Section Method). Then, MuCST learns the transcriptional features of spots by discriminating the attributed network and random one generated with permutation of gene expression profiles, where neighbors of spots are also in close proximity to each other in the transcriptional feature space.

On the data reconstruction issue, MuCST considers the morphological and transcriptional features of spots as complementary modalities, which fuses these heterogeneous features of spots with multi-modal contrast learning (Fig. 1**C**). In detail, the morphological and transcriptional features of spots are projected into a shared subspace, where MuCST aligns the distributions of two types of features such that they are subjected to the identical distribution. In this case, the heterogeneity of spatially resolved data is removed at the feature level, facilitating the down-stream analysis. Then, MuCST restores histology image *I*^∗^and transcriptional profiles of spots *X*^∗^by minimizing the reconstruction errors, i.e., ∥ *I* - *I*^∗^ ∥ and ∥ *X* - *X*^∗^ ∥, thereby improving the compatibility of the reconstructed data, facilitating users for further study. Finally, MuCST employs principle component analysis (PCA) to independently learn the transcriptional and morphological features of spots from the reconstructed morphological and transcriptional information of spots, and fuses them for down-stream analysis. In experiments, we demonstrate that the fused features of spots facilitate the critical applications of spatially resolved data, including spatial domain identification, tumor micro-environment, and denoising spatial transcriptomics (Fig. 1**D**).

In all, MuCST is a flexible network-based model for integrating spatially resolved data, which jointly learns the compatible features of spots with multi-modality contrastive learning. It attempts to address heterogeneity of spatial omic data with the hierarchical strategy and network model, where heterogeneity of data is modeled and removed at feature level. Furthermore, MuCST not only facilitates users for downstream analysis, but also serves as the pre-processing step for spatially resolved data, such as denoising and restoring the original data.

### Benchmarking MuCST with simulated spatially resolved data

To evaluate the performance of MuCST, we first utilize simulated spatially resolved data, including the morphological information, spatial coordinate and transcription profiles of cells, in which the ground truth subpopulation assignment for each cell is known (Methods, Supplementary Section 1.3). The typical algorithms, such as CCA (with principle component analysis (PCA) for features) (40), AE (auto-encoder) (41), MUSE (33), as well as the concatenation of various features (feature concat.), and multi-modal integrative algorithms such as conST (35), ConGI (34), stMVC (31), and stMGATF (32), are selected as baselines. In details, CCA and ConGI learns features by maximizing cross-modality correlation, and AE integrates multi-modal data with a bottleneck layer to reconstruct the original data. conST directly concatenates heterogeneity multi-modal features as attributes of cell network, and stMVC and stMGATF employ semi-supervised strategy for integrative analysis of multi-modal data.We select adjusted Rand Index (ARI) (42) to measure performance of various algorithms for identifying cell sub-populations in simulated data.

By following MUSE (33), we first validate the capacity of algorithms to learn discriminative features from each modality. In detail, the number of sub-populations in the full multi-modal space is fixed by randomly merging a different group of clusters for each modality. As the number of clusters decreases, CCA and feature concatenation-based algorithms are similar to the single-modality approaches, whereas MUSE and MuCST are significantly superior to others, demonstrating that these algorithms capture the discriminative features of multi-modal data (Fig. 2**A**, Supplementary Fig. S1-2(**A**)). Furthermore, MuCST outperforms MUSE in all these cases. Specifically, ARI of MuCST is 0.880 ± 0.024 (for 15 clusters), 0.887 ± 0.020 (for 10 clusters), and 0.883 ± 0.018 (for 6 clusters) respectively, which is 2%∼5% higher than MUSE. Visualization of the latent features with t-SNE (43) for all these algorithms demonstrates that MuCST learns the compatible and discriminative features of cells that precisely characterize and model structure of cell sub-populations (Supplementary Fig. S1**A**, Fig. S2**C**). However, feature concatenation, AE and conST, are even worse than single-modality approaches, indicating that heterogeneity of modalities greatly affects performance of algorithms. Although ConGI neglects spatial information, resulting in an undesired performance. stMVC and stMGATF achieve an excellent performance since these algorithms take 50% labels as prior information, and performance of them also decreases with the number of clusters increases.

**Fig. 2.**
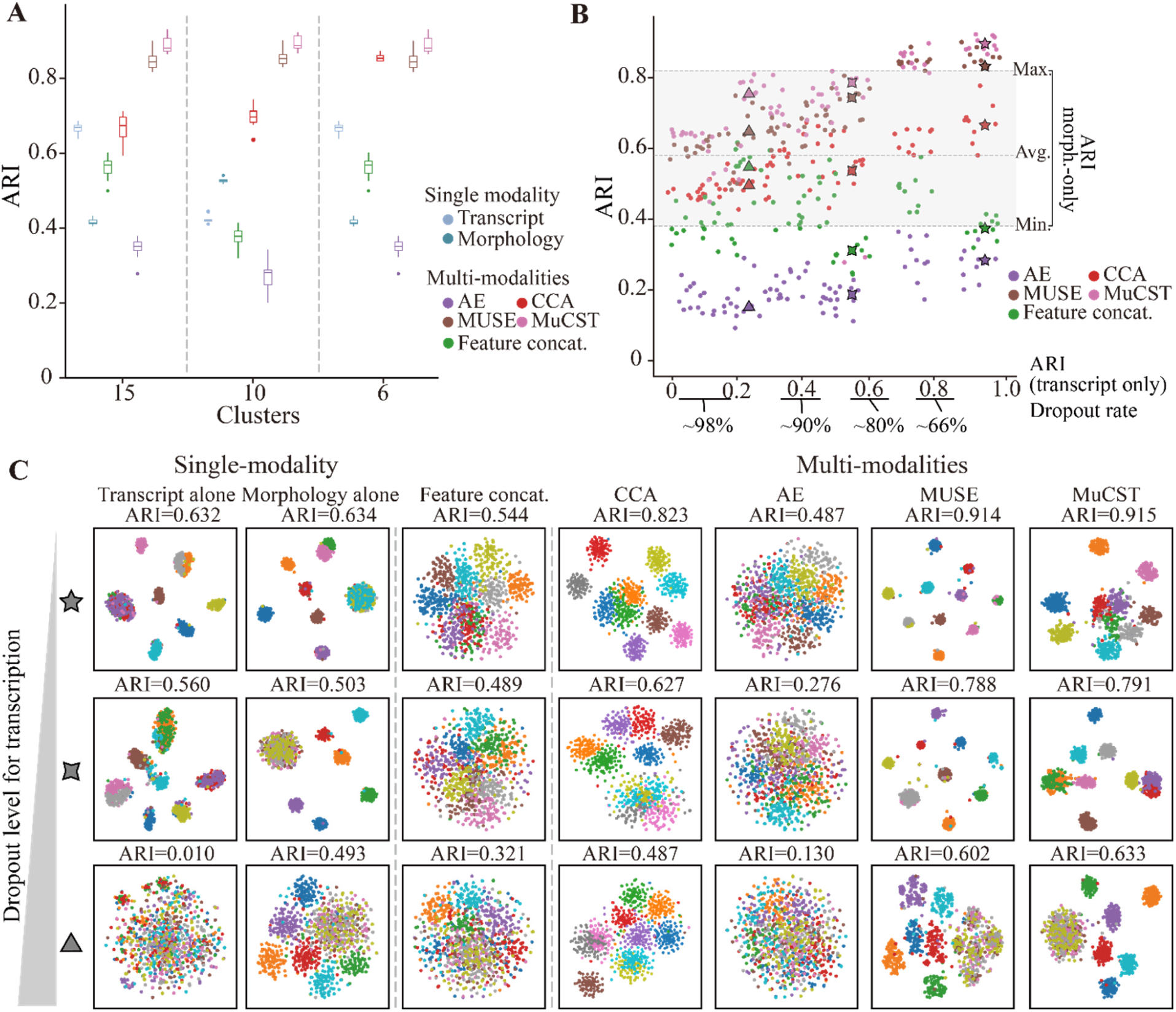
Performance of various algorithms on the simulated data. **(A)** ARI of identifying ground truth high-resolution subpopulations from lower-resolution single-modality subpopulations (k=15, 10 or 6), where 1,000 cells with transcriptional and morphological profiles are simulated. And, box plot is based on 10 replicates with median (center line), interquartile range (box) and data range (whiskers). **(B)** ARI of identifying ground truth clusters by varying the range of dropout levels from the transcriptional modality, where dashed lines denote minimum, average and maximum ARI of morphology modality alone, x axis represent ARI of PCA on transcript modality alone, y axis denotes ARI of combined-modality methods. **(C)** tSNE visualizations of latent representations from single and combined-modality methods, where ground truth subpopulation is labeled with various colors in simulation.

Next, we further validate performance of MuCST by degrading data quality of one modality, where two persistent strategies, i.e., dropouts and noise, are selected to perturb transcription data. By varying the level of transcript dropout rate, average ARI of morphology-alone method is 0.6 (Fig. 2**B** and Supplementary Fig. S2**B**, horizontal dashed lines). As the data quality of transcript degrades (i.e., the dropout rate increases, from right to left along with x-axis), performance of all these multi-modal methods drops dramatically. In all, MUSE, stMVC, stMGATF, and MuCST are much more robust than others since the worst performance of them are similar to that of morphology-alone method (Fig. 2**B** region between ‘min’ and ‘max’ of morphology-alone). Furthermore, MuCST is inferior to tMVC and stMGATF, but is superior to others for all the dropout rates. The reason is that tMVC and stMGATF make use of 50% labels, whereas MuCST is unsupervised learning. These results demonstrate that MuCST automatically discriminates the high- and low-quality modality, thereby improving performance of algorithms. Visualization the latent space learned by various algorithms demonstrates that MuCST precisely models structure of ten sub-populations regardless of dropout rate, proving that MuCST captures intrinsic structure of multimodal data (Fig. 2**C** and Supplementary Fig. S2**D**). Notice that multi-modal methods, such as CCA, conST, ConGI, and feature concatenation, are dramatically affected by degradation of data quality in any modality because these algorithms solely focus on deriving the relations among various modalities, thereby resulting in sensitivity to data perturbation.

Then, simulated data are simultaneously contaminated by noise for transcript and morphology modalities using additive Gaussian random noise with various variances. Performance of all algorithms drops as variance of noise increases, and MuCST achieves the best performance for all noise levels (Supplementary Fig. S1**B**). In detail, AE, CCA and feature concatenation are inferior to single modality approaches, failing to learn compatible features from noised heterogeneous multi-modal data, whereas MUSE and MuCST precisely characterize noise and learn discriminative features from noise multi-modal data. Furthermore, performance gap between MuCST and MUSE dramatically enlarges as the variance of noise increases from 0.1 to 2, demonstrating that MuCST is more precise and robust than MUSE. There two reasons explain why MuCST is superior to baselines. First, MuCST employs attributed cell network model for multi-modal data, which provides a better and comprehensive way to characterize intrinsic structure of multi-modal data by fully exploiting directed and indirect relations among cells. Second, MuCST makes use of contrast learning to remove heterogeneity of multi-modal data, thereby improving quality of features. Third, MuCST reconstructs the original multi-modal data with the compatible features of cells.

To fully check discriminative of features, we employ PheoGraph, Hierarchical and K-means to perform the identification of cell sub-populations for features of cells learned by various algorithms (Supplementary Fig. S1**C**). And, performance of various algorithms is consistent with that of the original methods, indicating that these algorithms are insensitive to the selection of clustering methods. Furthermore, MuCST achieves a good balance between efficiency and accuracy, where it saves 50% running time of MUSE with even higher performance (Supplementary Fig. S1**D**). Comprehensive parameter analysis demonstrates that MuCST is insensitive to values of parameters (Supplementary section 1.3, Supplementary Fig. S1**E**). These results on simulated data demonstrate that MuCST not only captures discriminative features in multi-modal data, but also effectively avoids being confounded by data quality of modalities.

### MuCST significantly enhances performance of identifying spatial domains for various tissues

Spatial domains play an critical role for investigating structure and functions of tissues (44), and many algorithms are devoted to this issue. Thus, we validate performance of MuCST for identifying spatial domains, where three spatially resolved datasets from normal tissues are selected, including the LIBD human dorsolateral prefrontal cortex (DLPFC) dataset (45), 10 × Visium dataset of mouse brain tissue, and the human intestine dataset (46). Ten state-of-the-art clustering algorithms, including the nonspatial method SCANPY (17), and spatial methods Giotto (20), stLearn (28), SEDR (24), BayesSpace (21), SpaGCN (29), STAGATE (22), SpatialPCA (26), DeepST (30), ConGI (34), conST (35), stMVC (31), Spatial-MGCN (27), stMGATF (32), and MUSE (33), are selected as baselines. And, adjusted rank index (ARI) (42) is selected to measure performance of algorithms if the truth ground spatial domains are known.

DLPFC dataset consists of 12 tissue slices obtained from human brain that are manually annotated as six layers of dorsolateral prefrontal cortex (Layer1∼Layer6) and white matter (WM) on the basis of morphology and gene markers (Fig. 3**A**). MuCST significantly outperforms baselines on the identification of spatial domains in slice 151673 with ARI 0.641, while that of the best baseline is 0.620 (Fig. 3**A**, Supplementary Fig. S5). MUSE fails to identifies spatial domains since it ignores spatial information, which is critical factor for spatial domains. Furthermore, MuCST and STAGATE precisely discriminate Layer 6 and WM, which cannot be identified by other baselines (Fig. 3**A**, Supplementary Fig. S5). These results demonstrate that MuCST captures discriminative features of spots for spatially resolve data. Performance of various algorithms for all 12 slices of DLPFC in terms of ARI is summarized in Fig. 3**B** and Supplementary Fig. S7**A**, where MuCST outperforms baselines except for stMVC. In details, ARI of MuCST is 0.584 ± 0.060, whereas that of Spatial-MGCN is 0.498 ± 0.097, conST 0.437 ±0.052, DeepST 0.501 ± 0.077, and BayesSpace 0.432 ± 0.104 (mean ± standard deviation, Fig. 3**B**). stLearn and SCANPY are inferior to others because these algorithms either utilize one of modalities, or fail to fully integrate multi-modalities, which is consistent with assertion in simulated data. Furthermore, MuCST is more robust than others since its variance is much less than baselines, demonstrating that MuCST learns discriminative features of spots for all slices (Supplementary Fig. S3-S6).

**Fig. 3.**
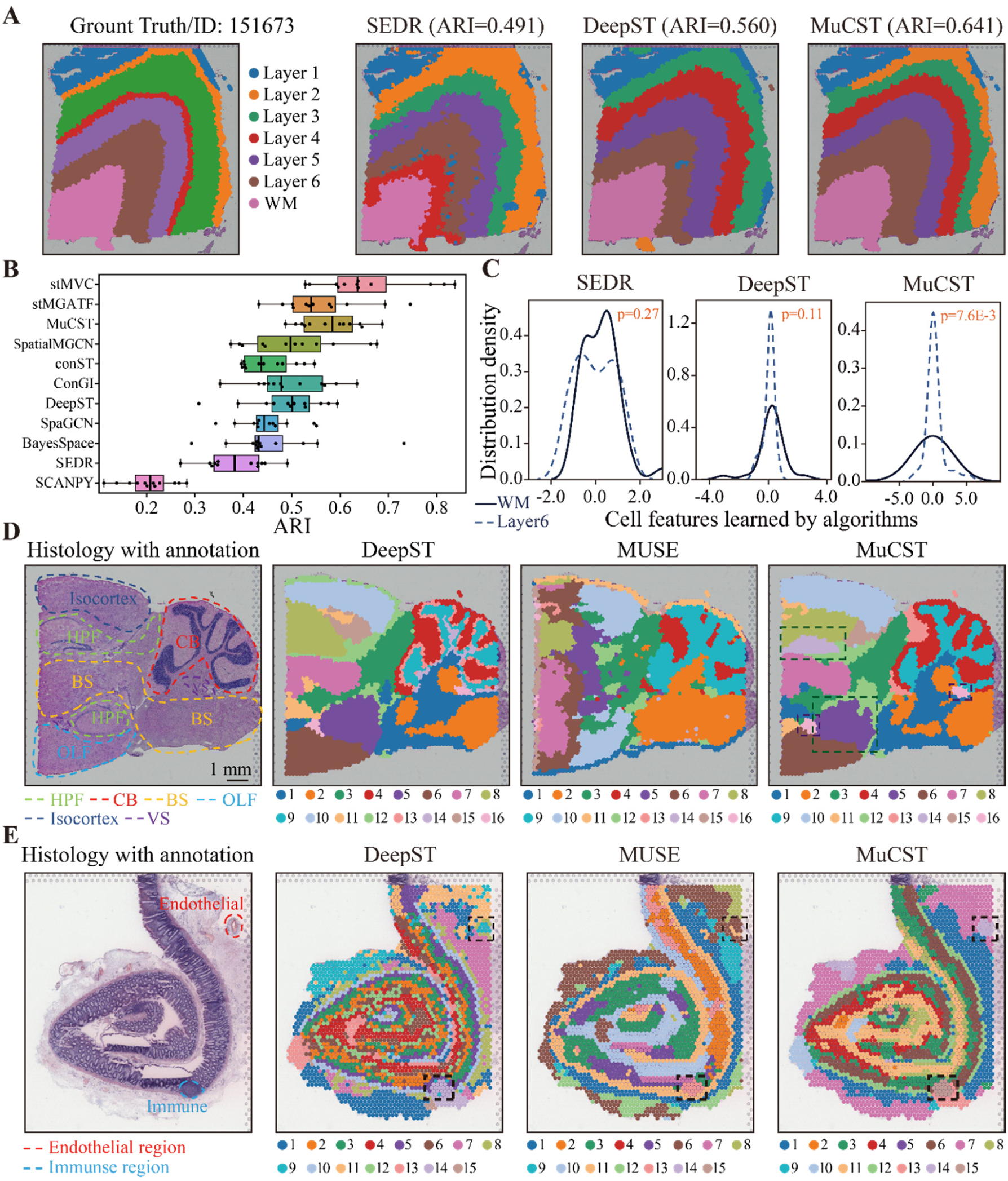
MuCST significantly enhances performance of spatial domain identification in normal tissues. **(A)** Ground truth segmentation of cortical layers and white matter (WM) in slice 151673 of DLPFC data, and visualization of spatial domains identified by SEDR, DeepST, and MuCST in slice 151673. **(B)** Boxplot of ARIs of various algorithms for spatial domains in all 12 slices of DLPFC, where x-axis denotes ARI, and the center line, box limits and whiskers denote the median, upper and lower quartiles, and 1.5× interquartile range, respectively. **(C)** Distribution density of cell features learned by SEDR, DeepST, and MuCST for Layer 6 and WM, where two side Kolmogorov–Smirnov test is employed for significance. **(D)** Annotated histology image of mouse brain posterior (left), and spatial domains obtained by different methods in delineating different structures of posterior brain. (E) Annotated histology image of human intestine (left), and spatial domains obtained by different methods in delineating spatial structures of human intestine.

Since MuCST integrates morphological and transcriptional features with contrast learning to remove heterogeneity of spatially resolved data, it is natural to validate quality of features. We compare distribution density of features learned by various algorithms for Layer 6 and WM (Fig. 3**B**). Surprisingly, features learned by MuCST significantly discriminate these two domains, whereas all these baselines fail to discriminate them (Fig. 3C, Supplementary Fig. S8). For example, deviation of distribution of MuCST is 1.69 and 7.62 for Layer 6 and WM respectively (p=7.6E-3, two-side Kolmogorov–Smirnov test), whereas that of MUSE is 0.56 (Layer 6) and 0.53 (WM) respectively (p=0.70, two-side Kolmogorov–Smirnov test). Furthermore, the single modality, such as transcript and morphology, also fails to discriminate Layer 6 and WM (Supplementary Fig. S5**A**, transcript: p=0.72, morphology: p=0.67, two-side Kolmogorov–Smirnov test). These results demonstrate that morphology is critical complementary information for characterizing and modeling spatial domains in spatial transcriptomics data, and MuCST precisely learns the discriminative and compatible spot features for spatial domains. Trajectory of spatial domains is fundamental for revealing mechanisms of biological evolution (47), and PAGA (48) is employed to infer relations of spatial domains identified by various algorithms, where the first spot in WM is selected as root cell. We further tested the robustness of MuCST by comparing the clustering accuracy with different hyper-parameters, and found that MuCST is insensitive to the encoder structure and the latent dimension (Supplementary Fig. S7**B**). MuCST precisely identifies the organization of the cortical layers derives from L1 to L6 and WM and achieves a PAGA score of 0.977, which is slightly lower than STAGATE’s 0.989, whereas other baselines mistakenly draw connections among various spatial domains (Supplementary Fig. S8**B**). These results demonstrate that MuCST accurately captures intrinsic structure and evolutionary relations of spatial domains.

Two additional 10x Visium datasets from mouse posterior (Supplementary Fig. S9(**A**)) and coronal brain (Supplementary Fig. S10(**A**)) are selected to further validate performance of MuCST, where the anatomical reference annotations are from the Allen Mouse Atlas. Fig. 3**D** visualizes annotation of posterior tissue, and spatial domains identified by various algorithms, where MuCST outperforms baselines (Supplementary Fig. S9(**B**)). Specifically, it is observed that single modality, such as transcript and morphology, accurately identifies Cerebellum (CB) area, but fails to distinguish Hippocampal Formation (HPF) and Brain Stem (BS) (Supplementary Fig. S9**A**). MUSE also fails to precisely identify spatial domains because it ignores spatial information, and DeepST cannot identify Cornu Ammonis (CA) and Dentate Gyrus (DG) areas because it employs the concatenation strategy to fuse the transcriptional and morphological features of spots. In contrast, MuCST offers flexibility in adjusting the importance of each modality, enabling clear identification of the CA and DG areas within the HPF areas in the mouse brain (surrounded by the dashed squares, Fig. 3**D**), as well as the Cerebellar Cortex and Dorsal Gyrus areas in the sagittal posterior mouse brain (surrounded by the dashed squares, Supplementary Fig. S9**B**), which are consistent with the reference annotations. Moreover, in the mouse coronal region, MuCST also effectively detects the Cornu Ammonis and Dentate Gyrus in HPF regions, demonstrating that MuCST delineates the spatial domain in more details (surrounded by the dashed squares, Supplementary Fig. S10(**B**). Furthermore, spatial domains identified by MuCST are with stronger regional continuity and fewer noise points than others. Silhouette Coefficient (SC) and Davies-Bouldin (DB) scores are employed to quantify quality of spatial domains, where MuCST has higher SC and lower DB score than baselines, demonstrating domains identified by MuCST are more precise from perspective of computation (Supplementary Fig. S9(**C**), Supplementary Fig. S10(**F**)). These results indicate that MuCST is also promising for characterizing complicated spatial domains in mouse brain.

We further adopt the mouse corona brain dataset with the histology image stained by antibodies (Alexa Fluor 488 anti-NeuN) and DAPI to validate the contribution of DAPI staining to MuCST, as shown in Supplementary Fig. S11(**A1**). The single transcript modality only identifies large spatial domains, failing to detect small ones, such as the “arrowlike” structure, i.e., dorsal gyrus areas within the hippocampus (Supplementary Fig. S11(**A2**), region surrounded by square with white border). From Supplementary Fig. S11(**A3**), it is easy to assert that morphology (DAPI staining) succeeds to identify the small spatial domains, whereas the identified spatial domains are irregular, which are inconsistent with annotation. MuCST precisely identifies the Ammon’s horn as well as the dentate gyrus structure in the hippocampus (Supplementary Fig. S11(**A4**), region surrounded by square with white border). And, MuCST precisely identifies the layer structures of the cortex region, where the obtained domains are coherent spatial structures, demonstrate that MuCST can effectively integrate morphological information extracted from DAPI staining images.

The human intestine dataset is further selected to verify performance of MuCST apart from brain slice, where four major spatial domains, such as epithelium, muscle, immune and endothelium region, are involved (Fig. 3**E**) (46). In the H&E image, endothelial and immune regions appear spatially grouped (rather than layered). Notice that MUSE and MuCST precisely identify these four spatial domains, whereas baselines fail to discriminate them (Fig. 3**E**, Supplementary Fig. S12**A**). Furthermore, morphology is much more precise than transcript, SEDR, SpaGCN, STAGATE, conST, ConGI, and Spatial-MGCN for identifying spatial domains in intestine dataset. The reason is that morphology dominates the intestine dataset, and current baselines fail to balance transcript and morphology, resulting in an undesirable performance. Interestingly, MuCST precisely identifies all these four spatial domains, which are highly consistent with annotation (Supplementary Fig. S12**B**-**E**). Furthermore, it also outperforms single-modality, demonstrating that MuCST reaches a good balance between transcript and morphology. These results show that MuCST enhances the identification of spatial domains for spatially resolved data from various tissues.

### MuCST significantly enhances performance of identifying spatial domains for various tissues

Spatial transcriptomics technologies are widely applied to disease research, and it is natural to investigate the generalization power of the proposed MuCST for revealing tumor heterogeneity from cancer spatially resolved data. Six public spatial transcriptomics datasets, including 10x Visium human breast cancer data, human primary pancreatic cancer (PDAC) data (49), human invasive ductal carcinoma (IDC) data (21), human HER2 breast cancer data (50), human prostate cancer data (51), and Alzheimer’s disease mouse brain slice datasets, are selected. The human breast cancer data consist of 3,798 spots and 36,601 genes, which is manually annotated by pathologists based on H&E images and spatial expression of the reported breast cancer maker genes (24), including 20 regions and 4 main morphotypes, i.e., ductal carcinoma in situ/lobular carcinoma in situ (DCIS/LCIS), healthy tissue (Healthy), invasive ductal carcinoma (IDC), and tumor edge regions (Fig. 4**A** left).

**Fig. 4.**
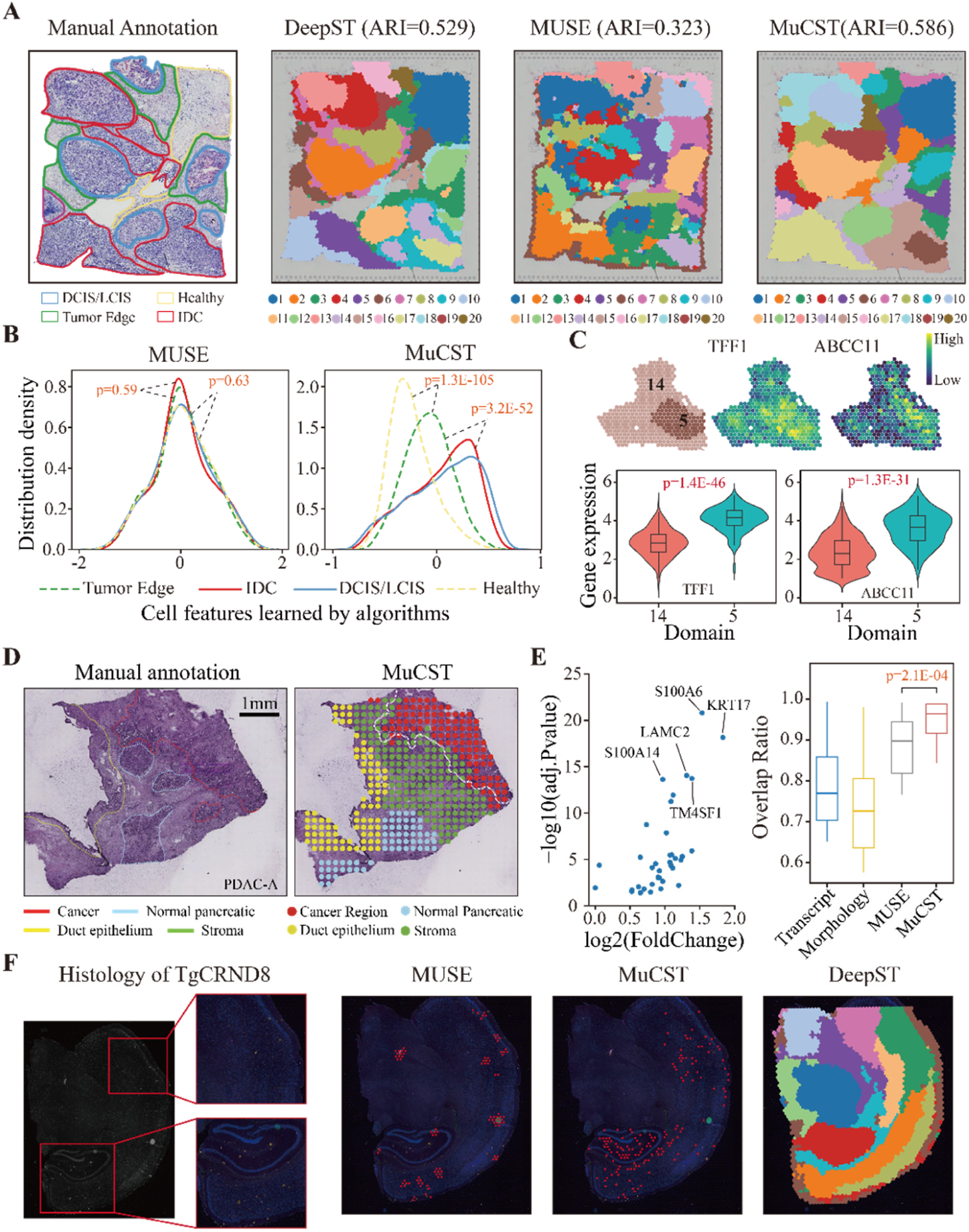
MuCST accurately identifies cancer-related domains for revealing tumor heterogeneity. **(A)** Visualization of annotation of human breast cancer data with healthy, tumor edge, IDC and DCIS/LCIS morphotype (left), and spatial domains identified by DeepST, MUSE and MuCST respectively (right). **(B)** Distribution density of spot features of breast cancer data learned by MUSE (left) and MuCST (right) for four morphotypes, respectively, where x-axis denotes features and y-axis denotes estimated distribution density (two-sided Kolmogorov– Smirnov test for significance). **(C)** Visualization of expression of TFF1 and ABCC11 between domain 5 and 14 (top), and violin plots of expression of these two genes (bottom). **(D)** Visualization of manual annotation of the human primary pancreatic cancer dataset with the normal, cancer, duct epithelium and stroma regions [49] (left), and spatial domains identified by MuCST (right). **(E)** Overexpressed genes in cancer regions through differential expression analysis between tumor and non-tumor regions characterized by MuCST (left), and overlap ratio of tumor region and marker genes identified by different algorithms (right). **(F)** Histology images of brain tissue sections from transgenic mice at the middle stage, where the zoomed in regions correspond to the primary region of amyloid plaque deposition (left). And, spatial domains identified by MUSE, MuCST and DeepST, respectively (right).

The spatial domains identified by various algorithms for human breast cancer are visualized in Fig. 4**B**, where MuCST outperforms all these baselines (Supplementary Fig. S13**A**). In details, cancer-related spatial domains identified by MuCST are highly consistent with the manual annotations (ARI=0.586, except for the semi-supervised algorithms, stMVC and stMGATF), whereas domains obtained by baselines with less regional continuity and more outliers (Supplementary Fig. S13**A**), implying that the proposed MuCST is also promising for characterizing and extracting cancer spatial domains. Furthermore, transcript alone and morphology alone are insufficient to fully characterize cancer-related spatial domains since ARI of them is 0.444 and 0.260 respectively (Supplementary Fig. S13**A**), indicating that morphological and transcriptional information complement for characterizing heterogeneity of tumors. Moreover, the 20 spatial domains identified by MuCST are divided into two categories with hierarchical clustering in terms of Pearson correlation coefficient, i.e., tumor and nontumor group, where the latter one includes spatial domains of tumor edge and healthy (Supplementary Fig. S13**C**). These results demonstrate that MuCST is also promising for characterizing and modeling tumor heterogeneity.

Then, we investigate whether spot features learned by various algorithms can characterize the tumor heterogeneity of breast cancers by discriminating these 4 major morphotypes. Fig. 4**B** describes the distribution density of spot features learned by MUSE (left) and MuCST (right), where MuCST precisely discriminate IDC, DCIS/LCIS, tumor edge and healthy morphotype. However, all these baselines, except for DeepST, failed to discriminate these four major morphotypes (Supplementary Fig. S13**B**), indicating the importance of spatial location and morphological information in revealing spatial domains within complex tissues. Interestingly, distribution density of spot features learned by MuCST not only precisely discriminates these major morphotypes, but also characterizes the evolutionary of morphotypes of breast cancer from healthy to tumor edge, and then to IDC (Fig. 4**B**, healthy: 0.67±1.25 vs tumor edge: 0.82±1.31, p=1.3E-105; tumor edge: 0.82±1.31 vs IDC: 0.94±1.38, p=3.2E-52, two-sided Kolmogorov–Smirnov test), which can be fulfilled by current baselines. These results demonstrate the proposed multi-modal contrastive learning strategy captures intrinsic structure of complicated cancer-related domains, providing an insight into mechanisms of tumors. To dissect tumor heterogeneity in tissue, differentially expressed genes (DEGs) among these four major morphotypes are obtained, which are highly associated with breast cancers. For example, *APOE* and *C1Q1* in tumor edge regions are associated with the differential abundance of tumor-associated infiltration of macrophages (TAM) that is critical for survival outcomes of patients due to its role in promoting tumor angiogenesis (52,53). And, the up-regulated genes are involved in immune and signal pathway, and down-regulated ones are associated with cell cycle process (Supplementary Fig. S13**D**). Moreover, heterogeneity of tumors results in hierarchical structure of spatial domains, i.e., IDC domain is further divided into two sub-domains (domain 5 and 14) (30). MuCST also precisely identifies them (Fig. 4**C**), where domain biomarker genes, such as *ABCC11* and *TFF1*, are differentially expressed. These results demonstrate that MuCST can reveal heterogeneity of tumors from various levels, i.e., from macro-level, such as spatial domains, to micro-level, such as feature and gene level.

To assess the performance of MuCST on dataset sequenced by legacy spatial transcriptomics rather than the 10x Visium, the human primary pancreatic cancer (PDAC) data is also adopted because of its high degree of heterogeneity, posing a great challenge on identifying spatial domains. The dataset is manually annotated, which consists of the normal, cancer, duct epithelium and stroma regions (49) (Fig. 4**D** left). Spatial domains identified by MuCST are highly consistent with the manually annotated areas (Fig. 4**D** right), whereas transcript alone, morphology alone, and baselines fail to identify pancreatic cancer related domains. Specifically, all these baselines mix the cancer and other domains (Supplementary Fig. S14**A**), which is avoided by MuCST, demonstrating its superiority of modeling tumor heterogeneity in complex tissues. We further conduct differential expression analysis between the cancer and normal areas (Fig. 4**E** left), and four of the top 5 DEGs, such as *KRT17*, *LAMC2*, *S100A14*, and *TM4SF1*, are bio-markers for PDAC (54, 55). Furthermore, these genes are significantly associated with survival time of patients, further substantiating the accuracy of spatial domains identified by MuCST (Supplementary Fig. S14**B**). The overlapping ratio of domain related DEGs identified by MuCST are more consistent with annotation than baselines (Fig. 4**E** right, two-tailed student test). These results demonstrate spatial domains facilitate the identification of pancreatic cancer bio-marker genes, which may provide clues for biologists for further study.

To validate the performance of MuCST on the identification of rare spatial domains in diseased related dataset, we employ an additional human HER2 breast cancer dataset generated by legacy spatial transcriptomics technology, where the zoomed in regions are manually annotated as rare spatial domains associated with breast cancer as shown in Supplementary Fig. S15 (**A**). Neither transcript nor morphology can identify these rare spatial domains, while MuCST precisely identifies these rare cancer domains that are surrounded by white solid squares. Furthermore, to overcome the low-resolution limitation of legacy platform, we use the super-resolution algorithm iStar (56) to validate the identified rare spatial domains, where the cancer epithelial cells are enriched in the identified small domains.

Furthermore, to ensure that MuCST is applicable to a variety of cancer spatial transcriptomics datasets, we further validate the performance of MuCST on human invasive ductal carcinoma (IDC) and human prostate cancer datasets, as shown in Supplementary Fig. S15(**B**). Spatial domains identified by MuCST are largely consistent with histopathological annotations, where 4 regions correspond to the annotated regions of predominantly IC (2, 6, 7, and 9), carcinoma in situ (5), benign hyperplasia (1), and predominantly non-tumor areas (3, 4 and 8, and 10). All these results demonstrate that MuCST precisely reveals the cancer-related spatial domains.

The additional 10x Genomics Alzheimer’s disease (AD) mouse brain dataset is selected because morphology dominates the data, where algorithms without morphology integration are invalid. Fig. 4**F** (left) visualize the immunofluorescence image of brain slice from a 5.7-month-old transgenic mouse, where the yellow amyloid plaques highlight the primary areas of amyloid-beta (Aβ) deposition. MUSE and MuCST identify discrete spatial domains associated with amyloid plaque accumulation, whereas baselines solely based on transcript only recognize coherent spatial domains of mouse brain tissue (Fig. 4**F** right, Supplementary Fig. S10**A**). Furthermore, MuCST identified the accumulation of amyloid plaques in the hippocampal region that cannot be accomplished by others. MuCST accurately identifies the plaque-covered areas while preserving the healthy brain regions on late-stage AD mouse brain slices (Supplementary Fig. S10**B**-**D**). We further conduct differential expression analysis between abnormal and normal spatial domains in the hippocampal region of mid-stage Alzheimer’s disease mouse brain slice. DEGs are involved in various Gene Ontology (GO) terms, such as gliogenesis and glial cell differentiation, which are associated with neurodegenerative diseases in brain regions (57) (Supplementary Fig. S10**E**). The spatial distribution of *Gfap* and *Cst3* are localized in the Aβ accumulation, which are highly expressed in AD brain slices and exhibit low expression in healthy brain slices, illustrating the potential effect of Aβ accumulation on gene expression (Supplementary Fig. S10**F**).

In summary, MuCST is more accurate to model and extract cancer-related domains, facilitating the understanding of tumor heterogeneity at various levels, which provides an alternative for integrative analysis of spatially resolved data.

### MuCST precisely characterizes and removes noise in spatially resolved data

Evidence demonstrates that spatially resolved data suffer from high noise levels because of dedicated procedures to preserve transcriptional and spatial information. Therefore, denoising is critical pre-processing for down-stream analysis (36, 58–60).

The proposed MuCST algorithm reconstructs the spatially resolved data with the compatible features, providing an alternative denoising strategy, and it is natural to investigate the capacity of MuCST for denoising. To validate the quality of reconstructed spatially resolved data, the third-party algorithm SCANPY is selected to perform spatial domain identifications on the original and restored data. Fig. 5**A** illustrates spatial domains identified by SCANPY by utilizing the original (left) and restored (right) transcript for slice 151673 of DLPFC, where ARI of algorithm dramatically improves from 0.181 to 0.480. Obviously, domains in the original data are full of noise and the boundary is unclear, whereas these domains in the reconstructed data are clearly and precisely identified. Specifically, WM and Layer 6 are clearly classified in the restored data, demonstrating that MuCST accurately captures intrinsic structure of spatially resolved data by removing noise. Furthermore, we compare distribution of features learned by SCANPY for WM and Layer 6 for the original and restored transcript, where difference between these two spatial domains is non-significant in the original data (Fig. 5**B** left), but it differs significantly (Fig. 5**B** right). In detail, the standard deviation of spot features from the original data is 1.03 and 0.76 for WM and Layer 6 respectively (p=0.72, two sample Kolmogorov-Smirnov test), whereas that of the restored data is 2.43 and 0.67 respectively (p=1.9E-79, two sample Kolmogorov-Smirnov test). These results show that MuCST precisely removes noise in spatially resolved data by exploiting the topological and multi-modality relations among spots, thereby enhancing the discriminative of spot features.

**Fig. 5.**
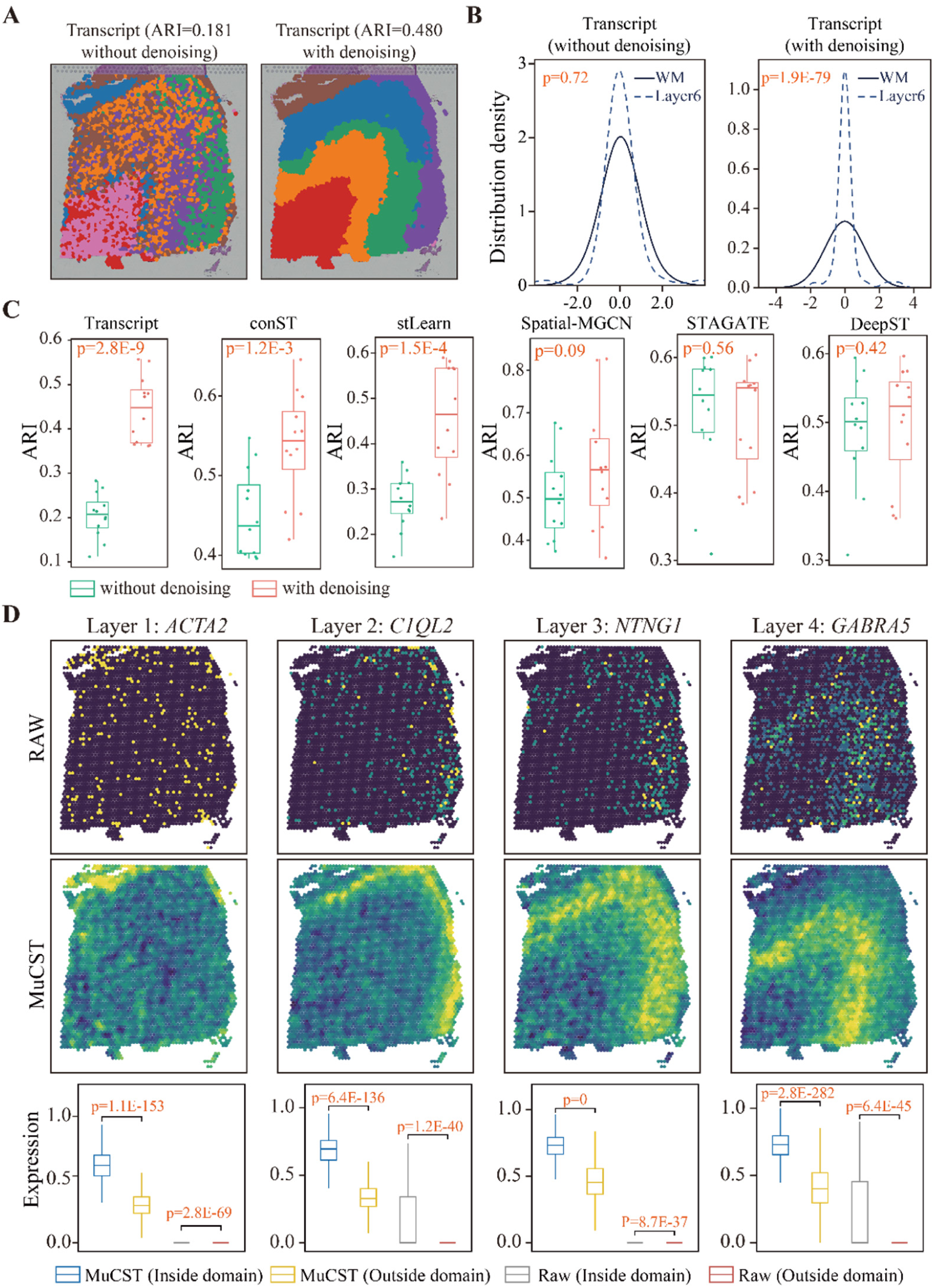
MuCST precisely removes noise in spatially resolved data to facilitate downstream analysis. **(A)** Visualization of spatial domains in slice 151673 of DLPFC identified by SCANPY based on the raw (left) and restored (right) spatial transcriptomics data, respectively. **(B)** Distribution density of spot features of slice 151673 learned by SCANPY for Layer 6 and WM from the original (left) and restored (right) spatial transcriptomics data respectively, where x-axis denotes features and y-axis denotes estimated distribution density (two-side Kolmogorov-Smirnov test for significance). **(C)** Distributions of ARIs of various algorithms for identifying spatial domains with the original and reconstructed DLPFC data respectively, where y-axis denotes ARI, and one-side student’s t test for significance. **(D)** Visualizations of the original (up), reconstructed data (middle) and expression of layer-marker genes (bottom) in slice 151673, where each column corresponds to one layer (two-side student’s t-test for significance).

To check whether the improvement of denoising is co-factored by algorithms and particular slice of DLPFC data, we apply all these baselines for spatial domain identification to the original and restored DLPFC data, where distributions of ARIs of all these algorithms on DLPFC data are described in Fig. 5**C**. Surprisingly, all these algorithms achieve a much better performance on the restored data than the original one, showing that the improvement of performance is not co-factored by the algorithms and particular slice. Moreover, SCANPY, stLearn and conST significantly improve performance of identifying spatial domains, and the other baselines also enhance performance with the restored data. For example, ARI of SCANPY (Transcript) increases from 0.205 ± 0.062 to 0.448 ± 0.071 (p=5.0E-9, one-side student’s t-test), whereas that of conST and stLearn soars from 0.271 ± 0.057and 0.437 ± 0.052 to 0.465 ± 0.118 and 0.544 ± 0.068 (p=1.3E-9, one-side student’s t-test). Even though improvement for Spatial-MGCN and DeepST is non-significant (Fig. 5**C**), the restored data lead to 14.98% and 4.6% improvement, respectively. The possible reason why STAGATE enlarges deviation of ARIs on the restored data is that it also performs denoising with an auto-encoder strategy, i.e., double denoising procedures result in an undesirable performance. These results demonstrate that multi-modality fusion is promising for characterizing and modeling noise in spatially resolved data, and MuCST can also serve as a pre-processing tool for down-stream analysis.

Since bio-marker genes are critical for spatial domains (46, 61), we then compare the expression of layer-marker genes for each layer between the original and restored data for slice 151673. Fig. 5**D** visualizes expression of bio-markers for each layer, such as *ACTA2* (Layer 1), *C1QL2* (Layer 2), *NTNG1* (Layer 4) and *GABRA5* (Layer 4) (45), for the original (up), and restored data (middle), where each column corresponds to a layer. It is easily observed that the bio-marker genes are not consistent with the structure of layers because of noise in the original data, while all these bio-marker genes are located in the corresponding domains. Then, we compare the expression of layer bio-marker genes within and outside of the corresponding layer in the restored data, where all these bio-marker genes are significantly expressed within domain than outsides (two-side student’s t-test, Fig. 5**D** bottom). Even though difference of these layer bio-marker genes is also significant, the difference is not as large as these in the restored data. These results demonstrate that MuCST precisely removes noise in spatially resolved data by augmenting expression of layer bio-marker genes, thereby improving quality of data.

The human breast cancer data is also selected validate performance of MuCST for denoising (Fig. 4**A**). We find that all these algorithms achieve higher accuracy on the restored data than the original one (Supplementary Fig. S17**A**). Particularly, transcript alone also enhances ARI from 0.444 to 0.546, and morphology alone increases ARI from 0.263 to 0.283, proving that improvement of accuracy is not co-factored by algorithms. Moreover, we also compare the distribution density of cell features learned by various algorithms for IDC and DCIS/LCIS, where the difference of features for these two layers is significant on the restored data, and is non-significant on the original one (Supplementary Fig. S17**B**, two sample Kolmogorov-Smirnov test), demonstrating that MuCST also precisely removes noise in breast cancer data. Finally, we also validate that the restored data also facilitate identification of spatial domains with bio-marker genes (Supplementary Fig. S17**C**).

In summary, MuCST precisely characterizes and removes noise with multi-modal contrastive learning, which can serve as critical pre-processing step for spatially resolved data.

### MuCST is applicable for spatial omics data with various platforms

Here, we investigate the applicability of MuCST with spatially resolved data generated with different platforms, such as STARmap (11), osmFISH (63), Slide-seq V2 (14), and Stereo-seq (15). The mouse primary visual cortex dataset obtained by STARmap (Fig. 6**A**) is selected, which is annotated as seven distinct layers from raw fluorescence data (Fig. 6**B**). And, the single modality transcript and morphology cannot fully characterize spatial domains with ARI 0.262 and 0.059 respectively, showing STARmap data is dominated by transcript. MuCST achieves the best performance with ARI 0.652, whereas that of STAGATE, SpaGCN, MUSE, ConGI, conST, stMVC, stMGATF is 0.586, 0.492, 0.057, 0.510, 0.400, 0.832 and 0.856, respectively (Fig. 6**A**, Supplementary Fig. S18**A**). The reason why MUSE fails to identify spatial domains is that it makes use of labels of spatial domains to guide feature learning, which is invalid if the balance of transcript and morphology loses. These results demonstrate that MuCST is also promising for integrating spatially resolved data generated with STARmap platform.

**Fig. 6.**
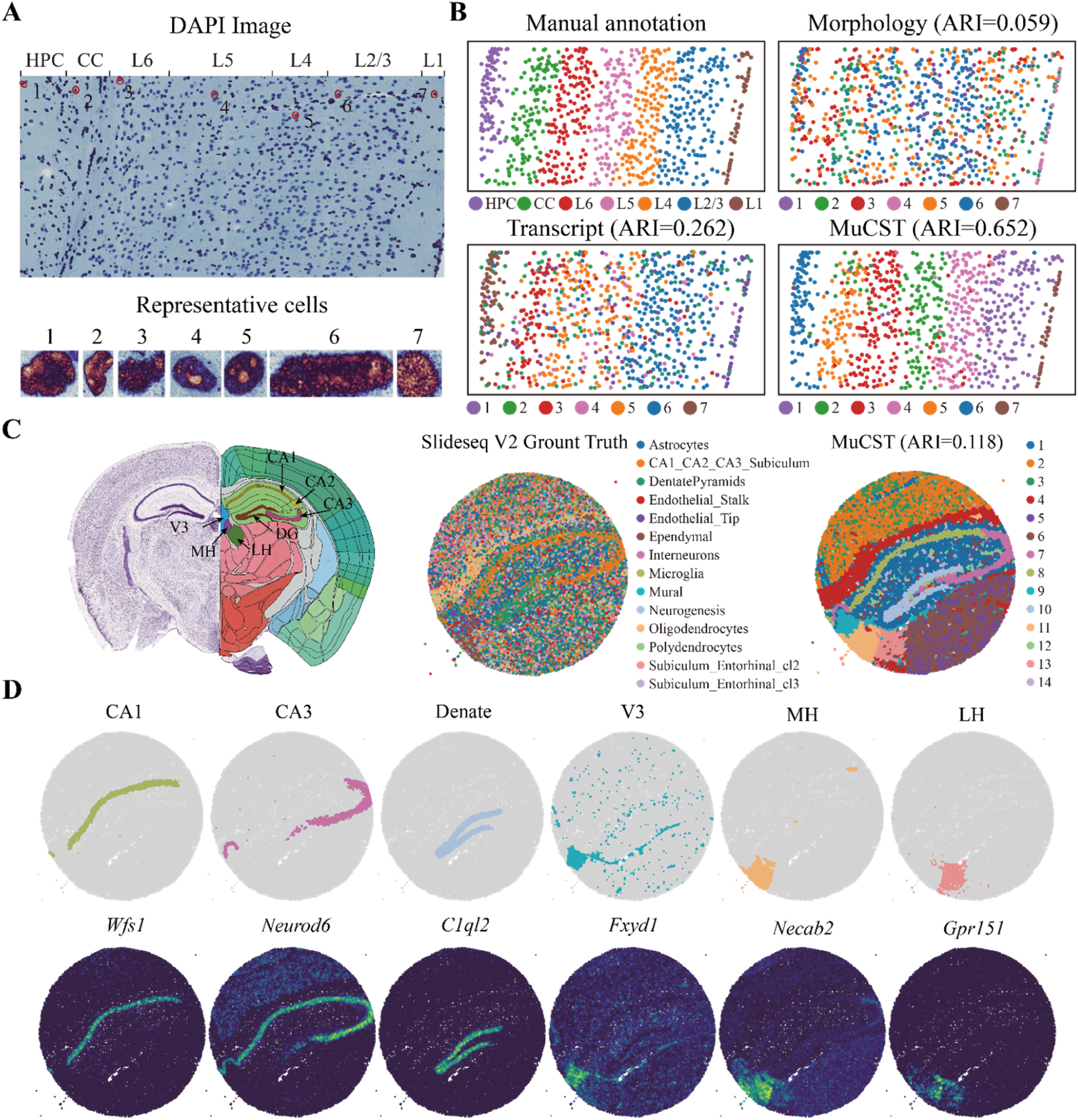
MuCST is applicable for spatially resolved data with various platforms. **(A)** Raw DAPI image of the V1 tissue annotated with seven functionally distinct layers (up panel), and seven representative cells from different layers (bottom panel). **(B)** Manual annotation of seven layers, and spatial domains identified by single modality and MuCST. **(C)** Visualization of mouse hippocampus tissue from Allen Mouse Brain Atlas (left), visualization of mouse hippocampus tissue annotated by (62) (middle), and spatial domains identified by MuCST (right). **(D)** Visualization of the spatial domains identified by MuCST, and the corresponding marker spatial gene expressions. The identified domains are aligned with the annotated hippocampus region of the Allen Mouse Brain Atlas.

We then validate the applicability of MuCST with three additional datasets from various platforms without morphological information, including the mouse brain cortex data with osmFISH (63), mouse hippocampus tissue with Slide-seq V2 (14), and mouse olfactory bulb tissue with Stereo-seq (15). The mouse brain cortex is a non-lattice shaped spatially resolved transcriptomics dataset generated by osmFISH, where spatial domains are labeled with different colors (Supplementary Fig. S18**B**). Compared to the state-of-the-art algorithms, MuCST and STAGATE achieve the best performance with ARI 0.500, demonstrating that the proposed model also works well for osmFISH data. The mouse hippocampus dataset generated with Slide-seq V2 are annotated based on the Allen Brain Atlas (62) (Fig. 6**C** left and middle panel). MuCST successfully identifies annotated spatial domains, including the dentate gyrus (DG) and the pyramidal layers within Ammon horn, which are further separated into fields CA1, CA2, and CA3. And, it outperforms DeepST on the delineation of CA3 and DG (Supplementary Fig. S18**C**). Moreover, the spatial distribution of expression of domain bio-marker genes are consistent with annotation of hippocampus regions, where each column corresponds to a domain identified by MuCST (Fig. 6**D**). In contrast, baselines mix some domains, for example, DeepST merges the MH and LH (Supplementary Fig. S18**C**).

Finally, we apply MuCST to the coronal mouse olfactory bulb tissue acquired with Stereo-seq (15), which is annotated with the DAPI image, including the olfactory nerve layer (ONL), glomerular layer (GL), external plexiform layer (EPL), mitral cell layer (MCL), internal plexiform layer (IPL), granule cell layer (GCL), and rostral migratory stream (RMS) (Supplementary Fig. S18**D**). SCANPY, DeepST, and MuCST precisely identify domains in the outer layers, i.e., ONL, GL, and EPL, while SCANPY mixes GCL with the outer IPL region in inner structure (Supplementary Fig. S18**E**). We further adopt the spatial distribution of marker genes of each anatomical region to validate MuCST-identified domains, where a well match is observed, demonstrating domains identified by MuCST are consistent with annotations (Supplementary Fig. S18**F**). Overall, MuCST effectively leverage the whole transcript and spatial information to discern the relevant anatomical regions.

## Discussion

The spatial transcriptomics measures gene expression at the cell level while retaining the associated spatial context, and the combination of gene expression, spatial coordinates, and morphological information of cells facilitates the identification of coherent cell patterns to understand the structure, functions and organization of tissues. However, it is highly non-trivial to integrate morphology and spatial transcriptomics because of noise and heterogeneity of data. In this study, we propose a novel multimodal contrastive learning algorithm (MuCST), which address the integrative analysis problem with joint learning.

In spatially resolved transcriptomics data, characterization of cellular heterogeneity is critical for revealing the structure and functions of tissues in health and disease. We demonstrate that MuCST precisely reveals subpopulation structure missed by either modality or compensate for noise (Fig. 2). Furthermore, we validate that MuCST also precisely reveals tumor heterogeneity from cancer spatially resolved data (Fig. 4). MuCST also accurately models noise of spatially resolved data under the guidance of feature learning, which can serve as pre-processing step for down-stream analysis (Fig. 5). Spatial domain identification is also prominent task for spatially resolved data, which is also another perspective to characterize tumor and tissue micro-environments. The multi-modal data provide a comprehensive way to characterize structure of tissues, which cannot be fulfilled with single modality. In this study, we show that MuCST precisely identifies spatial domains from the normal (Fig. 3) as well as tumor tissues (Fig. 4), covering different species, and diseases. We also validate that MuCST is also applicable to spatial omic data generated with various platforms (Fig. 6).

We see ample improvement space for MuCST from perspectives of data and methodology. Specifically, how to extend MuCST such that it can be applied to spatially resolved data generated with highly resolution technologies. Furthermore, compared to bulk-based sequencing data, the spatially resolved data are limited. How to combine the traditional bulk sequencing data with spatial omic data, i.e., increasing the number of modalities, is also interesting.

## Materials and Methods

### Data pre-processing and network construction

For all datasets, spots (cells) outside the main tissue regions are removed. Histology image is split into patches for each spot according to spatial coordinates, and morphological features of patches are learned with Resnet-50 (38) (denoted by ***m****_i_*). K-nearest neighborhood (KNN) is utilized to construct the cell network *G* = (*V*, *E*) with Euclidean distance of spatial coordinates of cells (*k* = 6 according to Ref. (62)). Then, weight on edge is calculated with similarity of morphological features of corresponding cells, i.e., weight *w_ij_* for the *i*-th and *j*-th cell is the cosine similarity of ***m****_i_* and ***m****_j_*. The raw expression profile of *n* cells *X̃* = [*x̃*_1_,…, *x̃_n_*] is normalized, log-transformed and scaled according to library size with SCANPY (17). By following Seurat (64), genes expressed in less than 10 cells are filtered. To overcome the low expression profile of cells, each cell is augmented with its neighbors. In details, given expression profile of the *i* -th cell *x̃_i_*, and its neighbors in *G*(*N_i_*(*G*) = *j*|(*i*, *j*) ∈ *E*}), the augmentation is performed by integrating morphological and expression of neighbors as

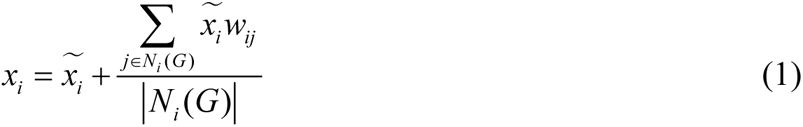

The attributed cell network 𝒢 = (*G*, *X*) is constructed by setting G as topological structure of cells, and the augmented expression profile of cells *X* as attributes of vertices. The random network 𝒢̂ = (*G*, *X̂*) of 𝒢 is generated by preserving the topological structure G and randomly permutating attributes of cells.

### Mathematical model for MuCST

MuCST consists of three major procedures, i.e., morphological feature learning, transcriptional feature learning, and multi-modal feature fusion. Therefore, the objective function of MuCST is also composed of three components corresponding to each of these procedures.

On the morphological feature learning issue, MuCST splits the histology image I into patches for each spot, and these patches are randomly noised. And, the pre-trained DNN model (38), followed by multilayer perception (MLP) (39) are adopted to learn the latent morphological features of spots. To enhance quality of features, SimCLR (37) is employed to discriminate the original and noised patches, and then MuCST fuses the morphological and transcriptional features of cells with multi-modal contrast learning. The histology image is reconstructed by minimizing the approximation, i.e.,

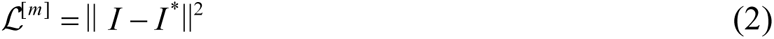

where *I*^∗^ is the reconstructed histology image.

On the transcriptional feature learning issue, MuCST employs graph neural network (GNN) to learn cell features by manipulating structure of attributed cell network 𝒢. Specifically, MuCST utilizes graph convolution network (GCN) (*ι* layers) (65) to learn cell transcriptional with structure as

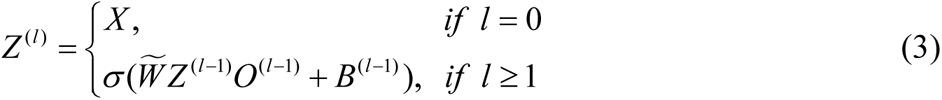

where 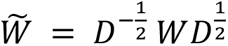 is the normalized adjacent matrix of 𝒢 by the degree sequencing matrix *D* (i.e., diagonal matrix with element as *d_ii_* = ∑*_j_*_=1_ *w_ij_*), *O* and *B* denote the trainable weight and bias matrix respectively, *σ*(⋅) is a non-linear activation function, *Z* is the latent transcriptional feature at the *l*-th layer. Then, MuCST employs GCN to reconstruct the expression profile of cells by using the learned transcriptional features *Z*^(*l*)^with structure as

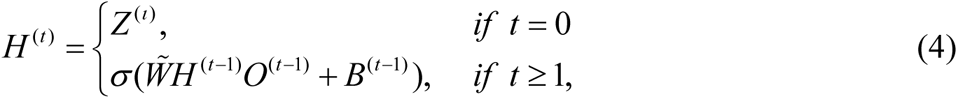

where *H*(*t*) denotes the reconstructed expression profiles at the *t*-th layer of GCN. Therefore, the loss function for expression reconstruction is defined as

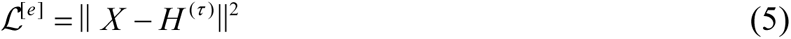

where parameter *τ* is the number of layers for decoder. To improve the discriminative of transcriptional features of cells, we expect features of cells belonging to the same spatial domains are close, whereas these from different domains apart from each other. For each cell *z_i_*, MuCST enforces it to be close to the center of its neighbors in the attributed cell network 𝒢 that is the mean of transcriptional features of them 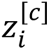, i.e., 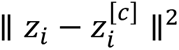. The random transcriptional features and pesudo center for the *i*-th cell (denoted as *ẑ_i_* and 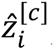) are obtained by applying GCN to random network 𝒢. Therefore, we maximize distance between *ẑ_i_* and 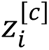 to discriminate cells belonging to various spatial domains, which is formulated as generative adversarial learning (66) as

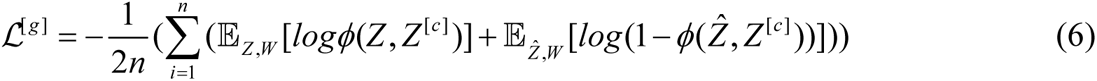

where *ϕ*(⋅) is the bilinear function, and *E*(*X*) denotes the mathematical expectation of matrix *X*, respectively.

On the multi-modal feature fusion issue, we employ a two-layer neural network 𝛶 to project the morphological and transcriptional features into a shared subspace to reduce the heterogeneity of cell features as

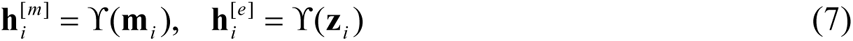

where the relations between morphology and spatial expression are implicitly exploited. To comprehensively fuse these two types of cell features, we expect the distributions of morphological and transcriptional features are consistent. Specifically, MuCST enforces the morphological features 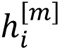 is close to the center of its neighbors for transcriptional features 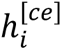, where can be fulfilled with contrastive learning (67) as

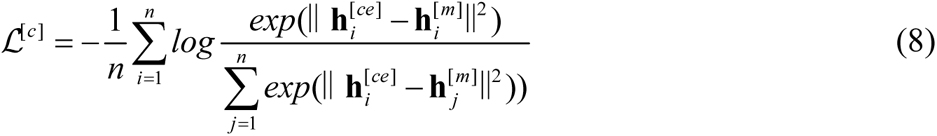

By combining Eq.(2), (5), (5), and (8), the overall objective function of MuCST is formulated as

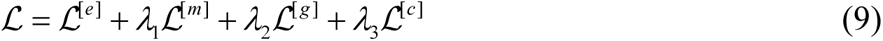

where parameter *λ*_1_, *λ*_2_, and *λ*_3_ control the relative importance of morphological information reconstruction, transcriptional feature consistence, and multi-modal fusion, respectively (The optimization and parameter setting are presented in Supplementary Section 1.2 and 1.3). And, in case when the histology image is absent, we set *λ*_1_ = *λ*_3_ = 0, e.t. the objective function of MuCST is reformulated as:

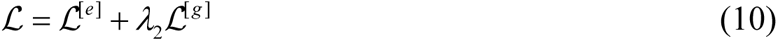

### Spatial domain identification

The morphological and transcriptional feature of cells, denoted by 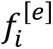 and 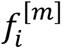, are obtained with PCA from the reconstructed morphological and transcriptional information, and are combined via a linear function as

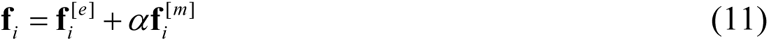

where parameter *⍺* controls the relative importance of morphological feature (here, we set *⍺* = *λ*_1_ in Eq. (10)). Then, MuCST identifies spatial domains by manipulating cell features *F* = [**f**_1_, …, **f**_**n**_]. Specifically, if the number of spatial domains is unknown, PhenoGraph (68) is employed to address this issue. Otherwise, Mclust (69) is adopted to identify spatial domains.

### Identification and functional analysis of differentially expressed genes

MuCST performs differential expression analysis of genes for each spatial domain by using Wilcoxon rank-sum test implemented in SCANPY package (17). Genes expressed in more than 80% of cells/spots in each domain, with a fold change ≥ 1 and an adjusted FDR ≤ 0.05, are selected as differentially expressed genes (DEGs). The filtered DEGs serve as input for gene ontology enrichment analysis, which is conducted with clusterProfiler (70). Enriched functional terms with -log10(adjusted P-value) are plotted.

### Clustering criteria

When manual annotation of spatial domains is absent, two extensively adopted clustering criteria, Silhouette Coefficient (SC) and Davies-Bouldin (DB) scores, are selected to validate the performance of clustering in terms of computation. Specifically, SC takes into account compactness within and separation across clusters as (b − a) / max(a, b), where a is the mean intra-cluster distance, and b is the mean nearest-cluster distance. It ranges between −1 and 1, where a higher score refers to more coherent clusters. SC=0 means that the sample is on or close to the boundary of neighboring clusters, whereas negative values denote potentially wrong clusters. DB score is the average ratio of within-cluster distances to between-cluster distances, favoring farther apart and less dispersed clusters with low values.

### Overlap ratio

In the dataset without ground truth, we employ the ratio of overlap between the spatial domain and marker genes as a metric to evaluate the performance of spatial domain identification. Specifically, we use the minimum expression value of the identified differential genes within the identified spatial domain as a threshold, then calculate the *R_gene_* for all spots with expression values exceeding this threshold and measure its overlap ratio with the *R_spatial_* of the identified spatial domain. The overlap ratio is formulated as:

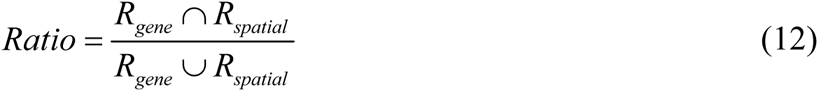

### Benckmarking

To comprehensively evaluate the performance of our method, we conducted several experiments to demonstrate its superiority over existing comparative methods. We compare MuCST with 10 state-of-the-art clustering algorithms, including the nonspatial method SCANPY (17), and spatial methods Giotto (20), stLearn (28), SEDR (24), BayesSpace (21), SpaGCN (29), STAGATE (22), SpatialPCA (26), DeepST (30), and one multi-modality method MUSE (33). All these algorithms are executed with the suggested values of parameters to achieve the best performance for fair comparison. For datasets with known annotations of spatial domains, the performance of algorithms is measured with adjusted rank index (ARI) (42) as

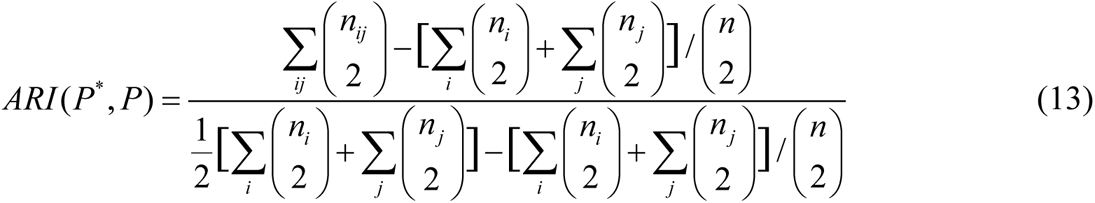

where *n* is the number of cells, *n_ij_* is the number of cells of class label *C*^∗^ ∈ *P*^∗^ assigned to cluster *C_i_* in partition *P*, and *n_i_*/*n_j_* is the number of cells in cluster *C_i_*/*C_j_* of partition *P*.

## Author contributions

Conceptualization: X.K.M.

Methodology: X.K.M., Y.W.

Investigation: Y.W.

Visualization: X.K.M., Y.W.

Supervision: X.K.M.

Writing—original draft: X.K.M., Y.W.

Writing—review & editing: X.K.M., Y.W.

## Data and materials availability

The code for the MuCST algorithm is implemented in Python and detailed tutorials are available at https://github.com/xkmaxidian/MuCST. The source code is released under the MIT License. All datasets analyzed in this paper are published datasets available for download. Human DLPFC is accessible within the SpatialLIBD (45) at http://spatial.libd.org/spatialLIBD. Human breast cancer, human invasive ductal carcinoma, Alzheimer’s disease mouse brain slices, and posterior and coronal mouse brain slices datasets are collected from the 10 × Genomics website at https://www.10xgenomics.com/resources/datasets. Human primary pancreatic cancer ST data (including gene expressions and H&E images) are downloaded from the Gene Expression Omnibus (GEO) database with accession number GSE111672. HER2-positive breast tumor ST data is accessible at https://github.com/almaan/her2st. Human prostate cancer ST data is hosted by the European Bioinformatics Institute (EBI), under the accession number EGAS0000100300. 10 × Visium human intestine spatial transcriptomics dataset are downloaded from the GEO database with accession number GSE158328. The mouse visual cortex STARmap data is accessible at https://www.wangxiaolab.org/data-portal-1. The mouse brain cortex osmFISH data is accessible at http://linnarssonlab.org/osmFISH. Slide-seqV2 dataset (14) is available at the Broad Institute Single Cell Portal at https://singlecell.broadinstitute.org/single_cell/study/SCP815. The processed Stereo-seq data (15) from mouse olfactory bulb tissue is accessible at https://github.com/JinmiaoChenLab/SEDR_analyses.

